# Furin Cleavage Site Is Key to SARS-CoV-2 Pathogenesis

**DOI:** 10.1101/2020.08.26.268854

**Authors:** Bryan A. Johnson, Xuping Xie, Birte Kalveram, Kumari G. Lokugamage, Antonio Muruato, Jing Zou, Xianwen Zhang, Terry Juelich, Jennifer K. Smith, Lihong Zhang, Nathen Bopp, Craig Schindewolf, Michelle Vu, Abigail Vanderheiden, Daniele Swetnam, Jessica A. Plante, Patricia Aguilar, Kenneth S. Plante, Benhur Lee, Scott C. Weaver, Mehul S. Suthar, Andrew L. Routh, Ping Ren, Zhiqiang Ku, Zhiqiang An, Kari Debbink, Pei Yong Shi, Alexander N. Freiberg, Vineet D. Menachery

**Author notes:** Equal contributions (Bryan A. Johnson, Xuping Xie). Co-senior authors (Pei Yong Shi, Alexander N. Freiberg, Vineet D. Menachery). **Corresponding Author:** Vineet D. Menachery **Address:** University of Texas Medical Branch, 301 University Blvd, Route #0610 Galveston, TX 77555 **Email:**.

## Abstract

SARS-CoV-2 has resulted in a global pandemic and shutdown economies around the world. Sequence analysis indicates that the novel coronavirus (CoV) has an insertion of a furin cleavage site (PRRAR) in its spike protein. Absent in other group 2B CoVs, the insertion may be a key factor in the replication and virulence of SARS-CoV-2. To explore this question, we generated a SARS-CoV-2 mutant lacking the furin cleavage site (ΔPRRA) in the spike protein. This mutant virus replicated with faster kinetics and improved fitness in Vero E6 cells. The mutant virus also had reduced spike protein processing as compared to wild-type SARS-CoV-2. In contrast, the ΔPRRA had reduced replication in Calu3 cells, a human respiratory cell line, and had attenuated disease in a hamster pathogenesis model. Despite the reduced disease, the ΔPRRA mutant offered robust protection from SARS-CoV-2 rechallenge. Importantly, plaque reduction neutralization tests (PRNT_50_) with COVID-19 patient sera and monoclonal antibodies against the receptor-binding domain found a shift, with the mutant virus resulting in consistently reduced PRNT_50_ titers. Together, these results demonstrate a critical role for the furin cleavage site insertion in SARS-CoV-2 replication and pathogenesis. In addition, these findings illustrate the importance of this insertion in evaluating neutralization and other downstream SARS-CoV-2 assays.

**Importance:** As COVID-19 has impacted the world, understanding how SARS-CoV-2 replicates and causes virulence offers potential pathways to disrupt its disease. By removing the furin cleavage site, we demonstrate the importance of this insertion to SARS-CoV-2 replication and pathogenesis. In addition, the findings with Vero cells indicate the likelihood of cell culture adaptations in virus stocks that can influence reagent generation and interpretation of a wide range of data including neutralization and drug efficacy. Overall, our work highlights the importance of this key motif in SARS-CoV-2 infection and pathogenesis.

**Article Summary:** A deletion of the furin cleavage site in SARS-CoV-2 amplifies replication in Vero cells, but attenuates replication in respiratory cells and pathogenesis in vivo. Loss of the furin site also reduces susceptibility to neutralization *in vitro*.

## Introduction

The rapid emergence of severe acute respiratory syndrome 2 coronavirus (SARS-CoV-2) at the end of 2019 ushered in a pandemic that has led to over 24 million cases and over 800,000 deaths ^1,2^. The novel coronavirus, like its predecessors severe acute respiratory syndrome coronavirus SARS-CoV and Middle East Respiratory Syndrome (MERS)-CoV induces potentially severe respiratory disease including fever, breathing difficulty, bilateral lung infiltration, and in many cases, death ^3,4^. While SARS-CoV-2 shares a similar genomic structure and protein homology with SARS-CoV, its ability to spread asymptomatically and cause a range of mild to severe disease distinguishes it from the earlier pandemic CoV ^5^. In exploring the differences, attention has been paid to the spike (S) protein, a key glycoprotein responsible for receptor binding and entry into the cell. Following receptor recognition, the S protein is subsequently cleaved at two sites, S1/S2 and the S2’ site to facilitate virus entry into a cell. Initial structural work has indicated that SARS-CoV-2 has greater affinity for the ACE2 receptor than the original SARS-CoV ^6,7^. In addition, changes in the N-terminal domain in addition to the receptor-binding domain indicate potential differences in attachment that may drive changes to transmission or virulence ^8^. However, the majority of attention has focused on a potentially critical insertion of a furin cleavage site upstream of the S1 cleavage site in spike ^9^. Absent in other group 2B CoVs, the four additional amino acids (PRRA) form the classic RXXR motif cleaved by many serine proteases when added to the conserved R found at the SARS-CoV-2 S1 cleavage site (PRRAR) ^10^. Notably, furin cleavage sites have been observed in other virulent pathogens like HIV, avian influenza strains (H5 and H7) as well as Ebola ^11^. In fact, furin cleavage sites are found in a number of other CoV family members including MERS-CoV, HKU1-Cov, and OC43-CoV ^12,13^; given the range of disease associated with these CoV strains, the furin cleavage site does not necessarily predetermine virulence. However, given its absence in other group 2B CoVs and the major differences in disease compared to SARS-CoV, a better understanding of the role of the furin cleavage site during SARS-CoV-2 infection is needed.

In this manuscript, we utilized a reverse genetic system to generate a SARS-CoV-2 mutant that lacked the furin cleavage site insertion ^14^. The mutant, ΔPRRA, had augmented replication and improved fitness in Vero E6 cells relative to wild type (WT) SARS-CoV-2. It also had reduced spike processing as compared to the WT virus. In contrast, the ΔPRRA mutant was attenuated in Calu3 cells, a human respiratory cell line, and had altered S protein processing as compared to Vero cells. *In vivo*, the ΔPRRA mutant had attenuated disease in hamsters despite robust, and sometimes augmented viral replication. Importantly, prior infection with the ΔPRRA mutant protected hamsters from subsequent rechallenge with WT SARS-CoV-2. Finally, neutralization assays that used the ΔPRRA mutant had lower PRNT_50_ values with both COVID-19 patient sera and monoclonal antibodies against the receptor-binding domain (RBD). Together, the results indicate a critical role for the furin cleavage site in SARS-CoV-2 infection and potential complications in interpreting research related to this virus infection.

## Results

The SARS-CoV-2 spike protein is >75% conserved in amino acid sequence across the group 2B CoV family with the majority of the differences occurring in the N-terminal domain and receptor-binding domain in the S1 portion (S. Fig. 1A). The insertion of a furin cleavage site (PRRA) upstream of the S1 cleavage site distinguishes SARS-CoV-2 from other group 2B CoV sequences including SARS-CoV and RATG13, the closest bat-derived CoV sequence (S. Fig. 1B). To evaluate the impact of the furin cleavage site insertion, we generated a mutant virus lacking the PRRA motif using our SARS-CoV-2 reverse genetic system (Fig. 1A) ^14^. Based on the SARS-CoV S protein structure, the insertion occurs in an exterior loop of the SARS-CoV-2 spike below the spike globular head and away from the receptor binding domain (Fig. 1B) ^15^. Using homology modeling based on the SARS-CoV S protein structure, we found the PRRA insertion in an extended loop (cyan). Deletion of PRRA insertion is predicted to shorten the loop, but importantly, not disrupt the overall structure of the spike protein. Following electroporation, we were able to recover the SARS-CoV-2 ΔPRRA mutant with stock virus titer roughly equivalent to the wild-type (WT) virus. Surprisingly, the SARS-CoV-2 ΔPRRA mutant virus produced a larger plaque size on Vero E6 cells than WT virus, suggesting potential changes in viral replication and spread in the absence of the furin cleavage site insertion (S. Fig. 1C).

**Figure 1.**
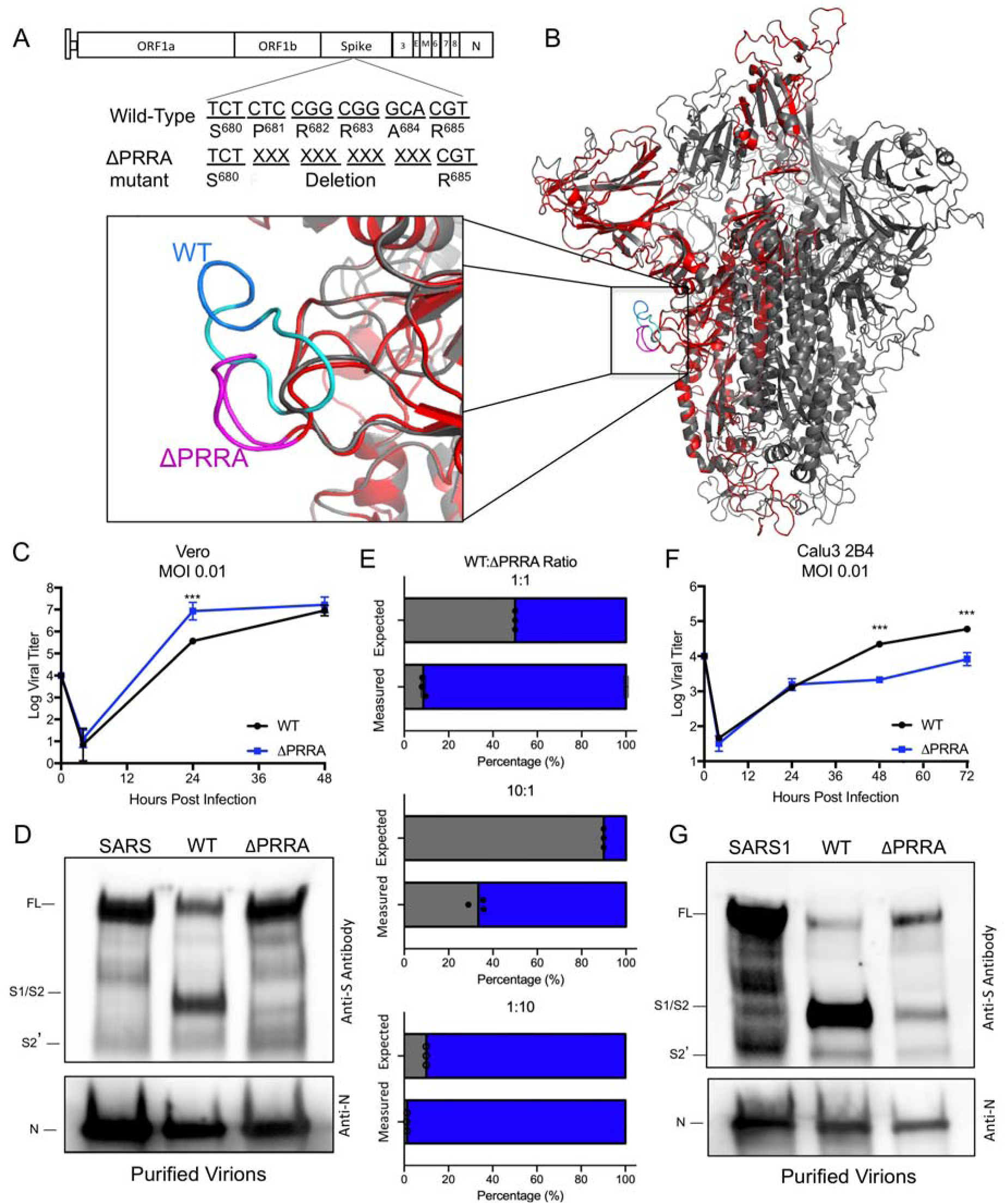
Distinct replication, spike cleavage, and competition for ΔPRRA. A) Generation of a SARS-CoV-2 mutant deleting the furin cleavage site insertion from the spike protein. B) Structure of the SARS-CoV-2 spike trimer with a focus on the furin cleavage site (inset). Modeled using the SARS-CoV-1 trimer structure (PDB 6ACD) (14), the WT SARS-CoV-2 trimer (grey) with SARS-CoV-2 PRRA deletion mutant monomer overlay (red). The loop (inset), which is unresolved on SARS-CoV-2 structures (AA 691-702), is shown in cyan on SARS-CoV-2 with the PRRA sequence in blue. The loop region in the PRRA deletion mutant is shown in pink. C) Viral titer from Vero E6 cells infected with WT SARS-CoV-2 (black) or ΔPRRA (blue) at MOI 0.01 (N=3). D) Purified SARS-CoV, SARS-CoV-2 WT, and ΔPRRA virions were probed with anti-spike or anti-nucleocapsid antibody. Full length (FL), S1/S2 cleavage form, and S2’ annotated. E) Competition assay between SARS-CoV-2 WT (black) and ΔPRRA (blue) showing RNA percentage based on quantitative RT-PCR at 50:50, 90:10, 10:90, 99:1, and 1:99 WT/ ΔPRRA ratio (N=3 per group). F) Viral titer from Calu3 2B4 cells infected with WT SARS-CoV-2 (black) or ΔPRRA (blue) at MOI 0.01 (N=3). G) Purified SARS-CoV, SARS-CoV-2 WT, and ΔPRRA virions were probed with anti-spike or anti-nucleocapsid antibody. Full length (FL), S1/S2 cleavage form, and S2’ annotated. P-values based on Student T-test and are marked as indicated: *<0.05 ***<0.001.

### Distinct replication kinetics and spike cleavage for the ΔPRRA mutant in Vero E6 cells

To evaluate viral replication, we infected Vero E6 cells with the WT and ΔPRRA SARS-CoV-2. Previous work has found robust replication of SARS-CoV-2 in Vero E6 cell and these cells are often used for propagation, including for inactivated vaccine production ^16^. Following low MOI (0.01 plaque forming units (PFU)/cell) infection, both WT and ΔPRRA SARS-CoV-2 mutant replicated to similar end-point titers (Fig. 1C). However, the ΔPRRA had a 25-fold increase in viral titer at 24 hours post infection (HPI) relative to WT. The increased replication was accompanied by more cytopathic effect (CPE) at 24 HPI; by 48 HPI, both mutant and WT levels had nearly 100% CPE. Together, the results suggested the loss of the furin cleavage site augmented replication in Vero E6 cells.

We next evaluated spike processing of the ΔPRRA mutant relative to WT SARS-CoV-2 as well as the original SARS-CoV. Vero E6 cells were infected at an MOI of ~0.1 for 24 h and purified virions were isolated from supernatants using ultracentrifugation and a sucrose cushion. The pelleted virus was subsequently examined for spike and nucleocapsid (N) protein levels by western blotting. Following SARS-CoV infection, the majority of the spike protein was observed in its full-length form (98.6%) (Fig. 1D, S. Fig. 2A), consistent with the absence of processing. In contrast, the WT SARS-CoV-2 virions had a significant reduction in full-length spike protein (40.4%). Instead, the most abundant form of the spike protein was the S1/S2 cleavage product (59.6%). Finally, the ΔPRRA mutant spike protein had mostly full-length spike (85.5%), similar to SARS-CoV, with only minimal processing to the S1/S2 cleavage form (14.5%). Given similar levels of viral N protein, these results illustrate the differences in processing between SARS-CoV and SARS-CoV-2. In addition, the data show that processing of the SARS-CoV-2 spike is driven primarily by the furin cleavage site following infection of Vero E6 cells.

**Figure 2.**
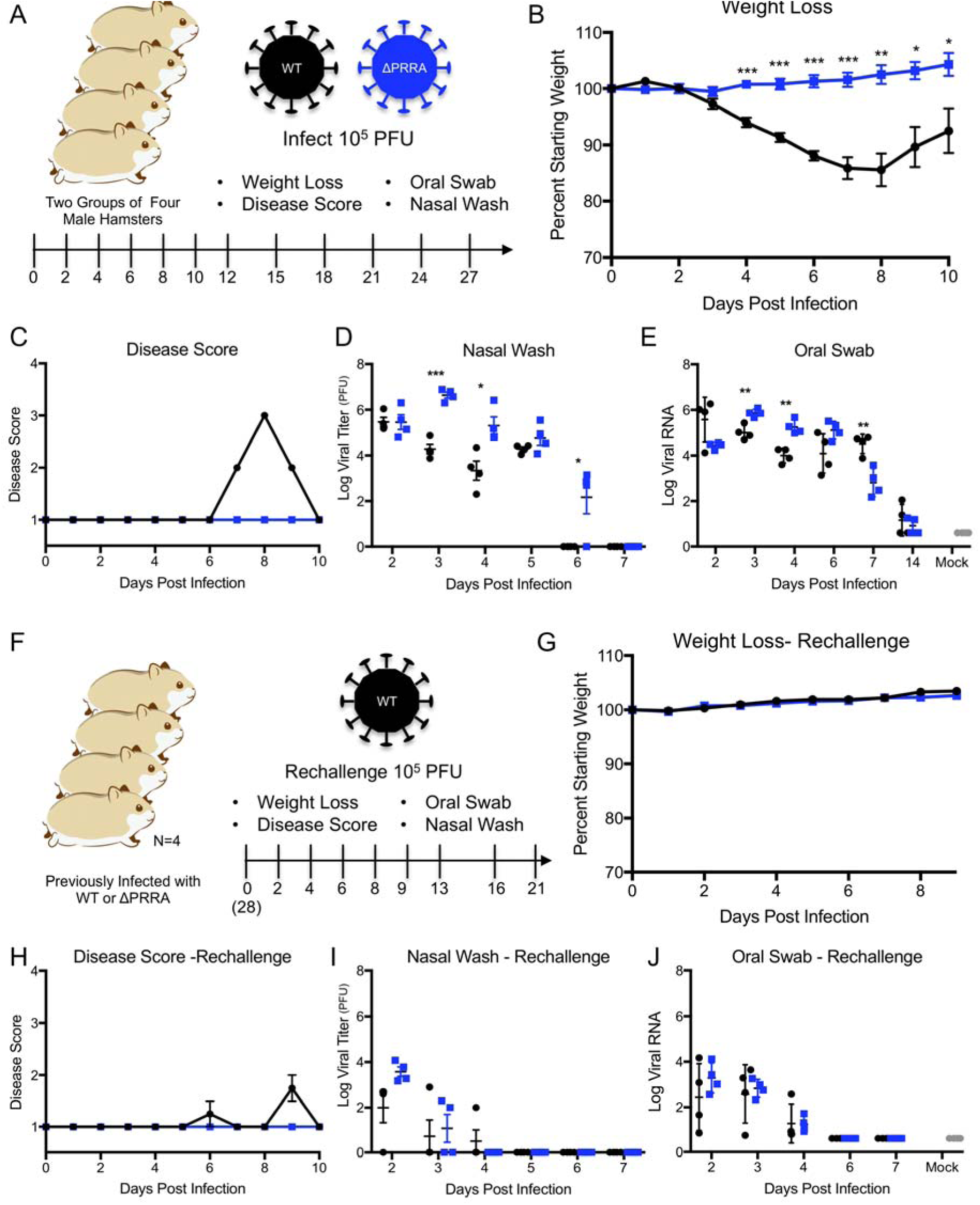
*In vivo* attenuation of ΔPRRA mutant. A) Primary SARS-CoV-2 challenge schematic. Two groups of male hamsters (N=4) were challenged with 10^5^ plaque forming units of either SARS-CoV-2 WT or ΔPRRA mutant and evaluated over a 28 day time course for B) weight loss, C) disease score, D) viral titer from nasal wash, and E) viral RNA from oral swabs. F) Schematic for rechallenge of previously infected hamsters. Twenty eight DPI, hamsters from SARS-CoV-2 WT and ΔPRRA were rechallenged with 10^5^ PFU of SARS-CoV-2 WT and evaluated for G) weight loss, H) disease score, I) viral titer from nasal wash, and E) viral RNA from oral swabs. P-values based on Student T-test and are marked as indicated: *<0.05 **<0.01 ***<0.001.

Given the replication advantage noted at 24 HPI (Fig. 1D), we next evaluated the fitness of the ΔPRRA mutant relative to WT SARS-CoV-2 in a competition assay. Using plaque forming units to determine the input, we mixed the WT and mutant viruses at different ratios in Vero E6 cells, and used a RT-PCR approach to evaluate their overall fitness after 24 hours (Fig. 1E, S. Fig 2B–C). At a 50:50 input ratio, the ΔPRRA mutant quickly outcompeted WT becoming nearly 90% of the viral population based on both RT-PCR distinguishing the two viruses. Similarly, a 90:10 WT to mutant input ratio resulted in ~65% of the viral sequences corresponding with the mutant virus, illustrating the advantage of the furin site deletion after only 24 HPI. The inverse 10:90 WT to mutant input ratio produced <3% of the WT virus and solidified the major advantage of the ΔPRRA mutant in Vero E6 cells. We further confirmed these data using deep sequencing analysis (S. Fig. 2D). Together, the results indicate that deletion of the furin cleavage site provides a fitness advantage in Vero E6 cells and a potentially potent cell culture adaptation.

### Attenuation of SARS-CoV-2 ΔPRRA in Calu3 respiratory cells

Having established augmented replication, altered spike processing, and enhanced fitness in Vero E6 cells, we next evaluated the ΔPRRA mutant in a more relevant cell type. Previously, Calu3 2B4 cells, a human lung adenocarcinoma cell line, had been sorted for ACE2 expression and used to study influenza and coronaviruses ^17,18^. In this study, we infected Calu3 2B4 with WT and ΔPRRA SARS-CoV-2 at MOI 0.01. Following infection, we found robust replication of WT SARS-CoV-2 peaking 72 HPI. In contrast, the ΔPRRA mutant virus was attenuated relative to WT beginning 48 HPI (Fig. 1F). At both 48 and 72 HPI, the ΔPRRA mutant had 1-log reduction in viral titer, contrasting the improved replication observed in Vero E6 cells. These data indicate that the loss of the furin cleavage site impairs replication in Calu3 cells and suggests SARS-CoV-2 requires the PRRA motif for efficient replication in these respiratory cells.

We subsequently repeated examination of the spike processing on virions from Calu3 2B4 cells for the ΔPRRA mutant. Similar to studies with Vero E6 cells, we infected Calu3 2B4 cells with SARS-CoV, SARS-CoV-2 WT, and ΔPRRA mutants at MOI 0.1. Given the reduced viral yields in Calu3 2B4 (Fig. 1F), we allowed replication to occur until 48 HPI before capturing supernatants and recovering purified virions. Consistent with findings in Vero E6 cells, the western blot of SARS-CoV virions showed the majority of spike protein was retained in the full length form (Fig. 1G, S. Fig. 2E). In contrast, the WT SARS-CoV-2 showed the majority of its spike protein had been cleaved to the S1/S2 form. Comparing to Vero E6 cells, spike processing to the S1/S2 form was more robust in Calu3 showing a ~87.3% at S1/S2 as compared to 59.6% in Vero E6 cells. Surprisingly, the ΔPRRA mutant also showed a significant increase in the S1/S2 cleavage product relative to its Vero spike blots. While the ΔPRRA has reduced overall infection, a clear band was visible at the S1/S2 cleavage site and represents more than 2 times as much S1/S2 cleavage product (33.1% vs 14.5%) compared to Vero E6 results. While more full-length spike is observed in the WT virus infection, the Calu3 results indicate that even without the furin cleavage site, there is significant processing of the SARS-CoV-2 spike. The results suggest that while the furin cleavage site is key in SARS-CoV-2 spike processing, other factors outside the PRRA insertion play a role in the efficient cleavage of the SARS-CoV-2 spike in a cell type-dependent manner.

### *In vivo* attenuation of Δ PRRA mutant

Having established contrasting results with *in vitro* studies, we next sought to evaluate the SARS-CoV-2 ΔPRRA mutant in an *in vivo* model. Early attempts found mouse models non-viable for SARS-CoV-2 infection ^19^; therefore we shifted to the hamster model which shows modest disease following infection with SARS-CoV-2 infection^20^. Four male hamsters were challenged with 10^5^ PFU of either WT SARS-CoV-2 or ΔPRRA mutant (Fig. 2A). The animals were subsequently monitored for 28 days with periodic measures of their body weight and disease signs. In addition, nasal washes and oral swabs were taken at day 2-7, 14, 21, and 28 days post infection (DPI). Following infection with WT SARS-CoV-2, hamsters steadily lost weight starting at day 2 and continuing through day 8 with peak weight loss nearing 15% (Fig. 2B, S. Fig 3A). These WT-infected hamsters also had disease scores that peaked between days 8 and 10, when animals showed signs including ruffled fur, hunched posture, and reduced activity requiring additional monitoring (Fig. 2C, S. Fig. 3B). Despite this severe disease, the WT-infected hamsters subsequently recovered and regained their starting weight by day 15 (S. Fig. 3A). In contrast, hamsters infected with SARS-CoV-2 ΔPRRA showed minimal weight loss over the course of infection (Fig. 3B, S. Fig. 3A). Over the first four days of infection, the ΔPRRA infected hamsters showed 2-3% weight loss, but remained close to their starting weight through day 10. In addition, the ΔPRRA mutant-infected hamsters had no change in disease score over the course of infection, distinguishing it from symptomatic disease observed following WT SARS-CoV-2 infection. The hamsters in both groups eventually gained a significant amount of weight after day 10 over the remainder of the 28-day time course (S. Fig. 3A).

**Figure 3.**
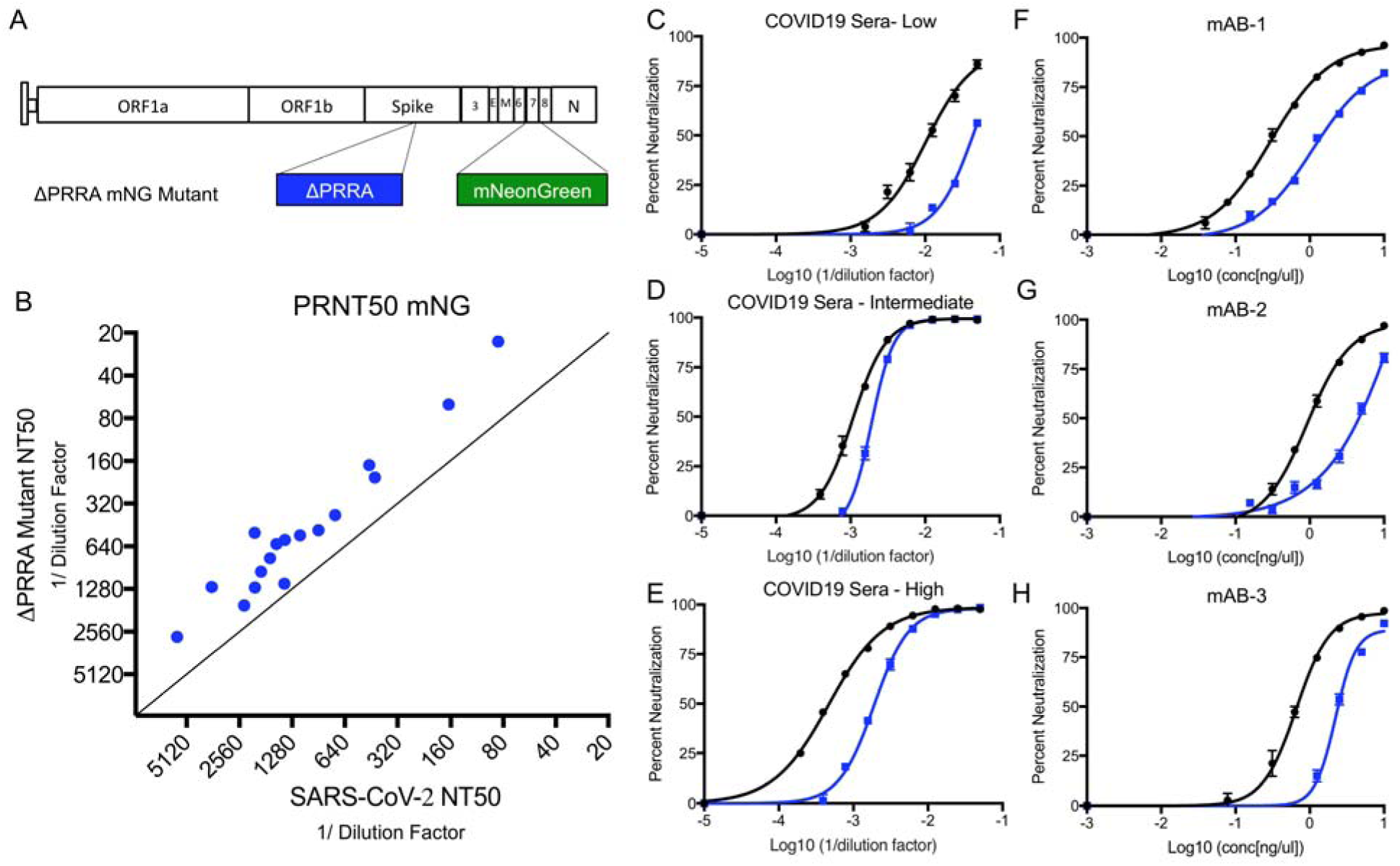
Antibody neutralization of ΔPRRA mutant. A) Schematic for SARS-CoV-2 ΔPRRA reporter virus expressing mNeonGreen (mNG) gene in place of ORF7 equivalent to previously described WT SARS-CoV-2 mNG virus ^21^. B) Plaque reduction neutralization (PRNT_50_) values as measured by changes to mNG expression. PRNT_50_ values plotted as Log (1/serum dilution) with ΔPRRA on Y axis and WT-SARS-CoV-2. C-E) Representative curves from C) low, D) intermediate, and E) high neutralizing COVID-19 patient sera. F-H) Neutralization curves from mAB-1 (F), mAB-2 (G), and mAB-3 (H), N=3.

Despite attenuated disease, the viral titers revealed augmented replication of the ΔPRRA mutant relative to WT SARS-CoV-2. Examining nasal washes, both WT and ΔPRRA infected hamsters had similar viral titers 2 DPI (Fig. 2D). However, augmented ΔPRRA replication was observed at both days 3 and 4 relative to the WT SARS-CoV-2. In addition, the WT virus was cleared from the nasal washes a day earlier than the ΔPRRA mutant, although no plaque forming units were detected after day 7 in either of the hamster groups. Evaluating oral swabs for viral RNA, a similar pattern was observed with augmented viral RNA at days 3 and 4 in the ΔPRRA mutant relative to WT (Fig. 2E). Notably, the viral RNA in the swabs stayed positive though day 7 with augmented WT viral RNA yield observed at the latest time point relative to ΔPRRA. Together, the results suggest that despite the attenuation in disease, the ΔPRRA mutant is capable of robust replication in the oral and nasal cavity of hamsters following infection.

### Infection with ΔPRRA mutant protects from SARS-CoV-2 rechallenge

We next evaluated if infection with ΔPRRA offered protection from further SARS-CoV-2 infection. Hamsters previously infected with either WT SARS-CoV-2 or the ΔPRRA mutant were rechallenged with 10^5^ PFU of WT SARS-CoV-2 28 days post initial infection (Fig. 2F). The rechallenged hamsters were monitored for disease and weight loss over a 21-day time course. Contrasting initial challenge, both WT and ΔPRRA infected hamsters were protected from weight loss following rechallenge (Fig. 2G and S. Fig. 3B). Hamsters infected with WT SARS-CoV-2 initially had no weight loss over the course of infection; however, mild disease (ruffled fur) was observed in a subset of animals (Fig. 2H). In contrast, rechallenge of ΔPRRA mutant-infected hamsters produced neither weight loss nor any evidence of disease. Nasal wash titers and viral RNA from oral swabs showed viral replication in both WT- and ΔPRRA-infected animals at day 2 and 3; however, the overall viral loads were significantly reduced compared to initial challenge (Fig. 2I & J). In addition, plaque forming virus appeared to be cleared around day 4 following rechallenge with low viral RNA loads found in oral swabs at corresponding time points. The results indicate that infection with the ΔPRRA mutant protects hamsters from disease upon rechallenge, but does not provide sterilizing immunity.

### Loss of furin cleavage site alters COVID19 serum neutralization

We next sought to evaluate the impact of the furin cleavage site deletion on virus neutralization by COVID-19 patient sera and monoclonal antibodies (mAB) against the SARS-CoV-2 receptor binding domain (RBD). To quantitate neutralization, we generated a ΔPRRA mutant containing the mNeonGreen (mNG) reporter in open reading frame 7 and subsequently compared neutralization results to the WT SARS-CoV-2 mNG reporter assay as previously described ^21^ (Fig. 3A). Examining seventeen COVID-19 human sera samples, we found nearly uniform reduction in PRNT_50_ values against the ΔPRRA mutant as compared to the WT control virus (Fig. 3B). The lower PRNT_50_ values were observed in sera whether the samples were from low, intermediate, or high neutralizing COVID19 patients (Fig. 3C–E) and averaged a 2.3-fold reduction across the 17 human sera. The consistency in the reduction suggested several possibilities: One is that virions themselves are altered in the conformation of the spike on the surface. In this situation, the processing or loss of spikes may permit access to more cryptic sites on the spike S2 or other regions allowing the WT virus to be more easily neutralized by non-receptor binding domain antibodies. A second possibility is that the loss of the furin cleavage site leaves more intact spike molecules on the virion surface requiring more antibodies to neutralize the ΔPRRA mutant than WT SARS-CoV-2. To explore this question, we examined the ΔPRRA mutant neutralization in the presence of three monoclonal antibodies (mAB) that target the SARS-CoV-2 receptor binding domains (RBD) (Fig. 3F–H). Each mAB targets a different site in the RBD, but each had similar reduction in the mAB serum neutralization levels between WT and ΔPRRA. The need for more mAB or polyclonal COVID patient sera suggest that ΔPRRA has more spike proteins that must be neutralized. Together, the results highlight significant differences in the neutralization between the WT and ΔPRRA SARS-CoV-2 mutants.

## Discussion

The loss of the furin cleavage site in the SARS-CoV-2 spike has a major impact on infection and pathogenesis. Using a reverse genetic system for the SARS-CoV-2 WA1 isolate, we generated a mutant virus that deleted the four amino acid insertion (ΔPRRA). The loss of the furin cleavage site resulted in reduced infection in Calu3 respiratory cells and ablated disease in the hamster pathogenesis model of SARS-CoV-2. Despite attenuated disease on initial infection, the ΔPRRA infected hamsters were protected from subsequent challenge with WT SARS-CoV-2 indicating induction of robust immunity. Together, the results highlight the importance of the furin cleavage site insertion to SARS-CoV-2 infection and pathogenesis.

Notably, despite attenuated disease *in vivo*, the ΔPRRA mutant had advantages over WT SARS-CoV-2 and may complicate research studies. The ΔPRRA mutant has a fitness advantage over the WT strain and dominated *in vitro* competition assays in Vero E6 compared to WT virus. Importantly, the furin site deletion has been reported in SARS-CoV-2 preparations ^22^ and given its fitness advantage in Vero E6 cells, can easily become the dominant tissue culture adaption in virus preparations. Coupled with *in vitro* and *in vivo* attenuation, efforts must be made to verify and evaluate stocks prior to critical studies. This also has implications for manufacturing inactivated COVID19 vaccine on Vero cells ^23^. Similarly, the shift in antibody neutralization values of the ΔPRRA virus indicates the possibility of inaccurate results if this mutation appears and distinguishes results from pseudotyped particles with and without this furin mutations ^24^. Fortunately, the ΔPRRA mutation will under represent the level of SARS-CoV-2 neutralization rather than overstating protection level; however, with potential vaccine and therapeutics decisions resting on these PRNT_50_ values, accuracy must be paramount. Together, the data highlight the importance of recognizing the mutation for future SARS-CoV-2 experimental analysis.

Biologically, the loss of the furin site shifts the processing of the spike in a cell type dependent manner. In Vero E6 cells, ΔPRRA significantly reduces cleavage to the S1/S2 form on the virion and the spike protein remains in the full-length conformation mirroring the results observed for SARS-CoV. In contrast, WT SARS-CoV-2 processes nearly 60% of its spike to S1/S2 indicating that most spike on the virion surface have been cleaved. Notably, the spike processing is distinct in Calu3 2B4 respiratory cells. While SARS-CoV virions remain uncleaved with little S1/S2 cleavage product, WT SARS-CoV-2 has increased processing from full-length to the S1/S2 cleavage product than what was observed in Vero E6 cells. Surprisingly, ΔPRRA in Calu3 cells also had a shift toward the S1/S2 fragment which is absent in Vero E6 cells. The results indicate that while the majority of the spike cleavage in SARS-CoV-2 is mediated by the furin cleavage site, there is more spike processing in SARS-CoV-2 even in its absence. With known serine protease differences between Vero E6 and Calu3 2B4 cells ^25^, the results suggest that spike processing varies based on cell type and may contribute to altered infection and pathogenesis *in vivo*.

In hamsters, the loss of the furin site attenuates SARS-CoV-2 induced disease, but does not ablate ΔPRRA virus replication. Following challenge, hamsters infected with the ΔPRRA had minimal change in weight loss over the first 10 DPI. In contrast, WT SARS-CoV-2 infected hamsters lost ~15% of their body weight and showed signs of disease (hunching, diminished movement, ruffled fur). While WT infected hamsters recovered, the absence of disease in the ΔPRRA infected hamsters indicates a key role for the furin cleavage site in virulence. Surprisingly, the ΔPRRA mutant was not attenuated in virus replication. At 3 and 4 DPI, the ΔPRRA had augmented titers as compared to control in nasal washes and increased viral RNA in the oral swabs. Similarly, the virus cleared one day later than WT SARS-CoV-2 during primary challenge. The results suggest that the reduced disease observed following ΔPRRA challenge was not a result of attenuated replication in these tissues.

Despite the lack of weight loss from initial challenge, ΔPRRA infected hamsters were protected from further WT SARS-CoV-2 infection. After 28 days, ΔPRRA and WT infected hamsters were rechallenged with WT SARS-CoV-2 and were protected from weight loss. While mild disease was observed in one of the WT SARS-coV-2 infected hamsters, the ΔPRRA infected hamsters showed no evidence of disease. However, low viral loads were observed in both the nasal washes and oral swabs from both groups, suggesting that the hamsters could foster a low level of infection after rechallenge. Yet, the virus was rapidly cleared and failed to induce disease in both groups, suggesting that adequate protection had been induced. Together, the results suggest that ΔPRRA mutant, despite attenuated disease, induces sufficient immunity to protect hamsters from further SARS-CoV-2 infection.

Overall, the data presented in this manuscript illustrate the critical role the furin cleavage site insertion in the spike protein plays in SARS-CoV-2 infection and pathogenesis. In its absence, the mutant ΔPRRA virus is attenuated in its ability to replicate in certain cell types and to cause disease *in vivo*. However, the results are complicated by augmented replication and fitness in Vero cells. Similarly, altered antibody neutralization profiles indicate a critical need to survey this mutation in analysis of SARS-CoV-2 treatments and vaccines moving forward. Together, the work highlights the critical nature of the furin cleavage site in understanding SARS-CoV-2 infection and pathogenesis.

## Methods

### Viruses and cells

The recombinant wild-type and mutant SARS-CoV-2 are based on the sequence of USA-WA1/2020 isolate provided by the World Reference Center for Emerging Viruses and Arboviruses (WRCEVA) and was originally obtained from the USA Centers of Disease Control as described ^16^. Wild-type and mutant SARS-CoV-2 as well as recombinant mouse-adapted recombinant SARS-CoV ^26^ were titrated and propagated on Vero E6 cells, grown in DMEM with 5% fetal bovine serum and 1% antibiotic/antimytotic (Gibco). Calu3 2B4 cells were grown in DMEM with 10% defined fetal bovine serum, 1% sodium pyruvate (Gibco), and 1% antibiotic/antimitotic (Gibco). Standard plaque assays were used for SARS-CoV and SARS-CoV-2 ^27,28^. All experiments involving infectious virus were conducted at the University of Texas Medical Branch (Galveston, TX) or Emory University (Atlanta, Georgia) in approved biosafety level 3 (BSL) laboratories with routine medical monitoring of staff.

### Construction of ΔPRRA Mutant Viruses

Both wild-type and mutant viruses were derived from the SARS-CoV-2 USA-WA1/2020 infectious clone as previously described ^14^. For ΔPRRA mutant construction, the mutation was introduced into a subclone puc57-CoV2-F6 by using overlap PCR with primers ΔPRRA-F (5-GACTAATTCTCGTAGTGTAGCTAGTCAATCCATC-3) and ΔPRRA-R (5-GACTAGCTACACTACGAGAATTAGTCTGAGTC-3). The resulted plasmid was validated by restriction enzyme digestion and Sanger sequencing. Thereafter, plasmids containing wild-type and mutant SARS-CoV-2 genome fragments were amplified and restricted. The SARS-CoV-2 genome fragments were purified and ligated *in vitro* to assemble the full-length cDNA according to the procedures described previously^14^. *In vitro* transcription reactions were then preformed to synthesize full-length genomic RNA. To recover the viruses, the RNA transcripts were electroporated into Vero E6 cells. The media from electroporated cells were harvested at 40-hour post-infection and served as seed stocks for subsequent experiments. Viral mutants were confirmed by sequence analysis prior to use. Synthetic construction of SARS-CoV-2 ΔPRRA mutant was approved by the University of Texas Medical Branch Institutional Biosafety Committee.

### *In Vitro* Infection

Viral replication in Vero E6 and Calu3 2B4 cells were performed as previously described ^29,30^. Briefly, cells were washed with PBS and inoculated with SARS-CoV or SARS-CoV-2 at a multiplicity of infection (MOI) 0.01 for 60 minutes at 37 °C. Following inoculation, cells were washed, and fresh media was added to signify time 0. Three or more biological replicates were harvested at each described time. No blinding was used in any sample collections, nor were samples randomized.

### Virion Purification and Western Blot

Vero E6 or Calu3-2B4 cells were infected with WT or PRRA mutant viruses at an MOI of 0.01. At 24/48 HPI, the culture media were collected and clarified by low speed spin. Virus particles in the media were subsequently pelleted by ultracentrifugation through a 20% sucrose cushion at 26,000 rpm for 3 h by using a Beckman SW28 rotor. For western blot analysis, protein lysates were prepared from the pellets using 2X Laemmli Sample buffer (Cat# 161-073, BioRad, Hersules, Ca). Relative viral protein levels were then determined by SDS-Page followed by western blot analysis as previously described ^16,31^. Briefly, sucrose purified SARS-CoV-1, SARS-CoV-2, and SARS-CoV-2 ΔPRRA inactivated by boiling in Laemelli Buffer. Samples were loaded in equal volumes into 4-20% Mini-PROTEAN TGX Gels (Biorad# 4561093) and electrophoresed by SDS-Page. Protein was then transferred to polyvinylidene difluoride (PVDF) membranes. Membranes were then blotted with SARS-CoV Spike (S) specific antibodies (Novus Biologicals #NB100-56576), followed by probing with horseradish peroxidase (HRP)-conjugated anti-rabbit antibody (Cell Signaling Technology #7074S) as a secondary. Blots were then stripped and re-probed with SARS-CoV Nucleocapsid (N) specific antibodies (provided as a kind gift from Dr. Shinji Makino) and the HRP-conjugated anti-rabbit secondary. In both cases, signal was developed by treating membranes with Clarity Western ECL substrate (Bio-Rad #1705060) imaging on a ChemiDoc MP System (Bio-Rad #12003154). Densitometry was performed using ImageLab 6.0.1 (Bio-Rad #12012931).

### Competition Assay and Real-Time PCR

For competition assays, ratios (50:50, 90:10, 10:90 WT/ ΔPRRA) were determined by plaque forming units derived from viral stocks. Vero cells were infected at MOI 0.1 (WTn+ ΔPRRA) as described above. RNA from cell lysates were collected using Trizol reagent (Invitrogen). RNA was then extracted from Triazol using the Direct-zol RNA Miniprep Plus kit (Zymo Research #R2072) per the manufacturer’s instruction. Extracted RNA was then converted to cDNA with the iScript cDNA Synthesis kit (BioRad #1708891). Quantitative real time PCR (qRT-PCR) was performed with the Luna Universal qPCR Master Mix (New England Biolabs #M3003) on a CFX Connect instrument (BioRad #1855200). For differentiation between wild type SARS-CoV-2 and SARS-CoV-2 ΔPRRA genomes in competition experiments, Primer 1 (Forward - AAT GTT TTT CAA ACA CGT GCA G and Primer 2 (Reverse - TAC ACT ACG TGC CCG CCG AGG) were used to detect wild type genomes only. For detecting total genomes, Primer 1 and Primer 3 (Reverse - GAA TTT TCT GCA CCA AGT GAC A) were used. 8-point standard curves (1−10^1^ to 1−10^8^ copies/ μL) were utilized to quantify the signal. A primer annealing temperature of 63°C was used for all assays.

For detection of viral RNA the nasal washes and oral swabs of SARS-CoV-2 and SARS-CoV-2 ΔPRRA infected hamsters, RNA extraction, cDNA synthesis, and qRT-PCR were performed as described above. For qRT-PCR, Primer 1 and Primer 3 were utilized for all hamster samples.

### Deep Sequencing Analysis

RNA libraries were prepared with 300ng of RNA using the Click-Seq protocol as previously described ^32^ using tiled primers cognate to the SARS-COV-2 genome (accession number NC_045512.2) and the TruSeq i7 LT adapter series and i5 hexamer adaptors containing a 12N unique molecular identifier (UMI). Libraries were sequenced on the Illumina MiSeq platform with MiSeq Reagent Kit v2. Raw data was de-multiplexed using TruSeq indexes using the MiSeq Reporter Software. Fastp v0.12. ^33^ was used to trim adapter sequences and low-quality reads (q<25), remove reads less than 40 nts in length, and copy UMI sequences onto the read name. Reads were aligned with bowtie using the best parameter and allowing for up to two mismatches. The alignment index was generated from a single fasta file, which contained two 600nt reference sequences spanning the PRRA locus (23603-23616) of the wildtype (accession number NC_045512.2) and ΔPRRA genomes. The alignments were sorted and indexed using Samtools v1.9 ^34^, PCR duplicates were removed using umi_tools ^35^. Coverage at each position was determined with the genomecov function in bedtools v2.25.0 ^36^.

### Plaque reduction neutralization titer assay

Neutralization assays were preformed using mNeonGreen SARS-CoV-2 reporter neutralization assay as previously described ^21^. Briefly, Vero E6 cells were plated black μCLEAR flat-bottom 96-well plate (Greiner Bio-one™). On following day, sera or monoclonal antibodies were serially diluted from 1/20 with nine 2-fold dilutions to the final dilution of 1/5120 and incubated with mNeonGreen SARS-CoV-2 or ΔPRRA expressing mNeonGreen at 37°C for 1 h. The virus-serum mixture was transferred to the Vero E6 cell plate with the final multiplicity of infection (MOI) of 0.5. After 20 hours, Hoechst 33342 Solution (400-fold diluted in Hank’s Balanced Salt Solution; Gibco) was added to stain cell nucleus, sealed with Breath-Easy sealing membrane (Diversified Biotech), incubated at 37°C for 20 min, and quantified for mNeonGreen fluorescence on Cytation™ 7 (BioTek). The raw images (2×2 montage) were acquired using 4× objective, processed, and stitched using the default setting. The total cells (indicated by nucleus staining) and mNeonGreen-positive cells were quantified for each well. Infection rates were determined by dividing the mNeonGreen-positive cell number to total cell number. Relative infection rates were obtained by normalizing the infection rates of serum-treated groups to those of non-serum-treated controls. The curves of the relative infection rates versus the serum dilutions (log_10_ values) were plotted using Prism 8 (GraphPad). A nonlinear regression method was used to determine the dilution fold that neutralized 50% of mNeonGreen fluorescence (NT50). Each serum was tested in duplicates.

### Phylogenetic Tree, Sequence Identity Heat Map, and Structural modeling

Heat maps were constructed from a set of representative group 2B coronaviruses by using alignment data paired with neighbor-joining phylogenetic trees built in Geneious (v.9.1.5) using the spike amino acid sequences derived the following accession numbers: QHU79204 (SARS-CoV-2 WA1), QHR63300.2 (RATG13), QND76034.1 (HKU3), AGZ48828.1 (WIV1), AGZ48806 (RsSHC014), ALK02457 (WIV16), and AYV99817.1(SARS-CoV Urbani). Sequence identity was visualized using EvolView (http://evolgenius.info/) and utilized SARS-CoV Co-V-2 WA1 as the reference sequence. Tree shows the degree of genetic similarity of SARS-CoV-2 and SARS-CoV across a selected group 2B coronaviruses. Structural models were generated using SWISS-Model ^37,38^ to generate homology models for SARS-CoV-2 spike with and without the furin cleavage site insertion based on the SARS-CoV-1 trimer structure (PDB 6ACD). Homology models were visualized and manipulated in MacPyMol (version 1.3).

### Animals and ethics statements

Male Syrian hamsters (7-8 weeks old, 86–127 g) were purchased from Envigo. All procedures were conducted under an animal protocol approved by the UTMB Institutional Animal Care and Use Committee and complied with USDA guidelines in an AAALAC-accredited lab. Work with infectious SARS-CoV-2 in hamsters was performed in the Galveston National Laboratory BSL-4 laboratory. Animals were housed in microisolator caging equipped with HEPA filters in the BSL-4 laboratories.

### Hamster Infection studies

Hamsters were challenged with 10^5^ PFU of WT-SARS-CoV-2 or SARS-CoV-2 ΔPRRA by intranasal inoculation (i.n.). Hamsters were observed daily for the development of clinical disease and body weights were taken every day for the first 10 days of the study, then every third day. For each manipulation (viral infection, retro-orbital bleeds, nasal wash, or oral swab), animals were anesthetized with isoflurane (Piramal, Bethlehem, PA).

### Statistical analysis

All statistical comparisons in this manuscript involved the comparison between 2 groups, SARS-CoV or SARS-CoV-2 infected groups under equivalent conditions. Thus, significant differences in viral titer were determined by the unpaired two-tailed students T-Test.

## Data Availability

The raw data that support the findings of this study are available from the corresponding author upon reasonable request.

## Acknowledgements

Research was supported by grants from NIA and NIAID of the NIH to (AI153602 and AG049042 to VDM AI142759, AI134907, AI145617, and UL1TR001439 to P-YS; R01AI123449 to AF and BL; R24AI120942 (WRCEVA) to SCW). Research was also supported by STARs Award provided by the University of Texas System to VDM and trainee funding provided by the McLaughlin Fellowship Fund at UTMB. P-YS was also supported by CDC grant for the Western Gulf Center of Excellence for Vector-Borne Diseases, and awards from the Sealy & Smith Foundation, Kleberg Foundation, John S. Dunn Foundation, Amon G. Carter Foundation, Gilson Longenbaugh Foundation, and Summerfield Robert Foundation.

## Author Contributions

Conceptualization, XX, BAJ, ALR, MS, ANF, P-YS, and VDM; Methodology, BAJ, XX, BK, KGL, DS, ALR, ANF, P-YS and VDM.; Investigation, BAJ, XX, BK, KGL, AM, JZ, XZ, TJ, JKS, LZ, CS, MV, AV, DS, NB, JAP, ALR, KD, and VDM.; Resources, KSP, SCW, MSS, PR, ZK, ZA, P-YS, ANF, VDM; Data Curation, BAJ, XX, BK, KGL, AV, DS, ALR, MSS, KD, P-YS, ANF, VDM.; Writing-Original Draft, VDM; Writing-Review & Editing, BAJ, XX, BL, PA, MSS, KD, ZK, ZA, P-YS, ANF, VDM.; Visualization, XX, BAJ, BK, KGL, NB, ANF, VDM; Supervision, PA, SCW, MSS, P-YS, ANF, VDM.; Funding Acquisition, PA, SCW, PY-S, ANF, VDM.

---

**S. Figure 1.**
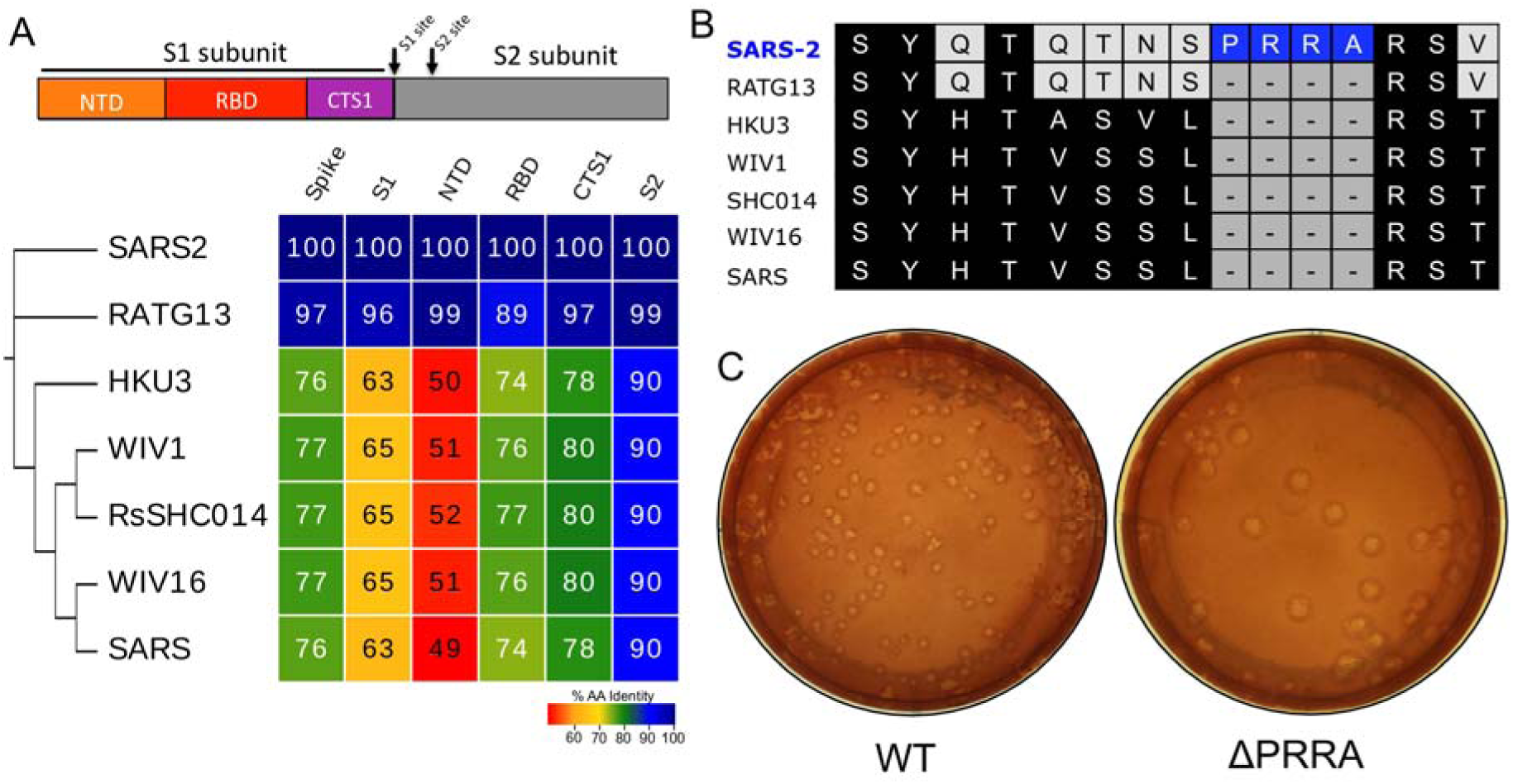
Furin cleavage site in SARS-CoV-2 spike. A) Diagram of the coronavirus spike protein domains and cleavage sites. The sequences of the indicated group 2B coronaviruses were aligned according to the bounds of total spike, S1, N-terminal domain (NTD), Receptor binding domain (RBD), and C-terminal of S1 (CTS1) and S2. Sequence identities were extracted from the alignments, and a heatmap of sequence identity was constructed using EvolView (http://www.evolgenius.info/evolview) with SARS-CoV-2 WA1 as the reference sequence. B) Alignment of the furin cleavage site of SARS-CoV-2 and the corresponding amino acids identities found closely related group 2B CoVs. The PRRA insertion is unique to SARS-CoV-2 C) Representative plaque morphology of WT and ΔPRRA SARS-CoV-2.

**S. Figure 2.**
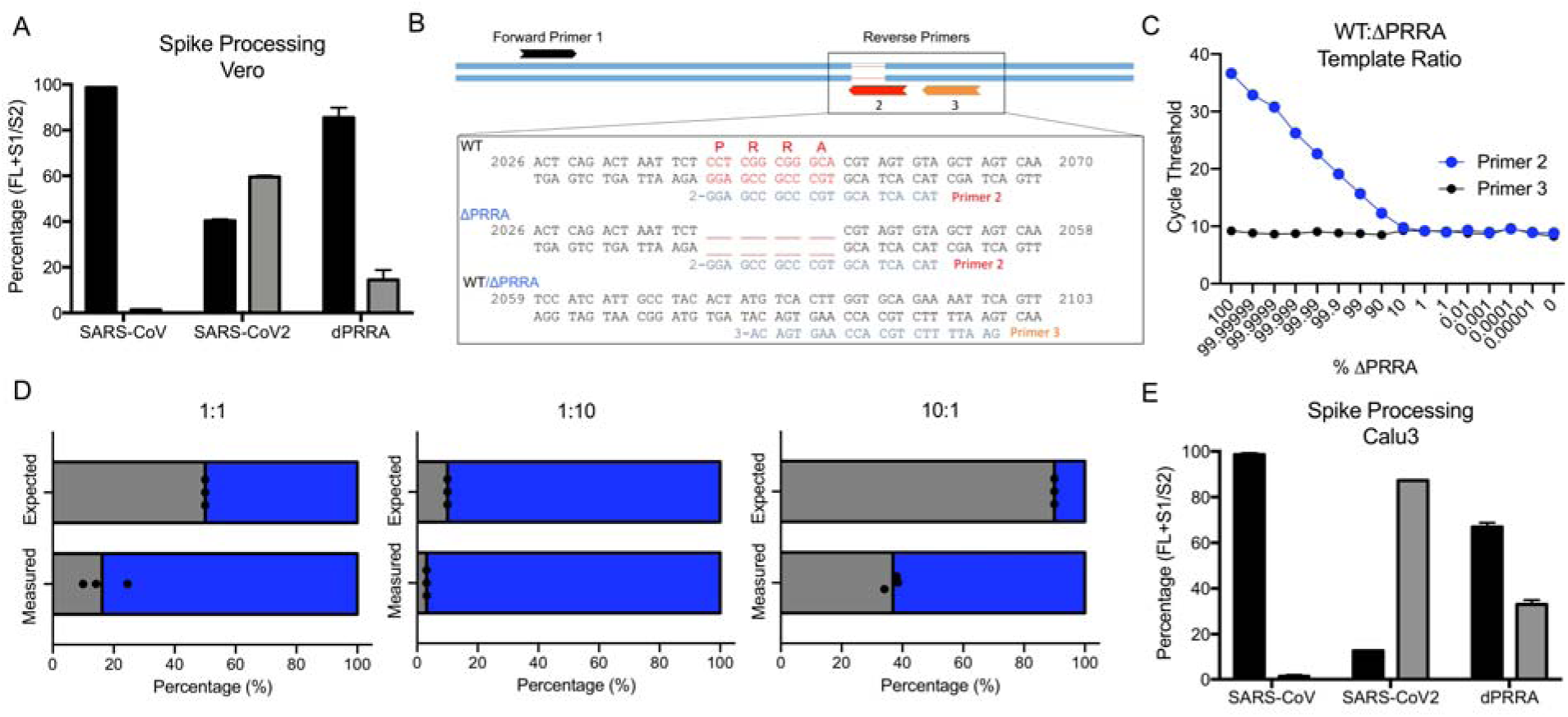
ΔPRRA mutant processing and competition with WT. A) Quantitation by densitometry of the full-length spike (Black) and S1/S2 cleavage form (Gray) from distinct western blot experiments in Vero E6 cells (n=2). B) Schematic of quantitative RT-PCR approach to detect deletion of the furin cleavage site. C) Primer curve validation with mixed WT to ΔPRRA plasmid ratio showing level of sensitivity. D) Deep sequencing results from ΔPRRA and WT competition assays based on percentage of total reads in that region (N=3). E) Quantitation by densitometry of the full-length spike (Black) and S1/S2 cleavage form (Gray) from distinct western blot experiments from Calu3 (n=2).

**S. Figure 3.**
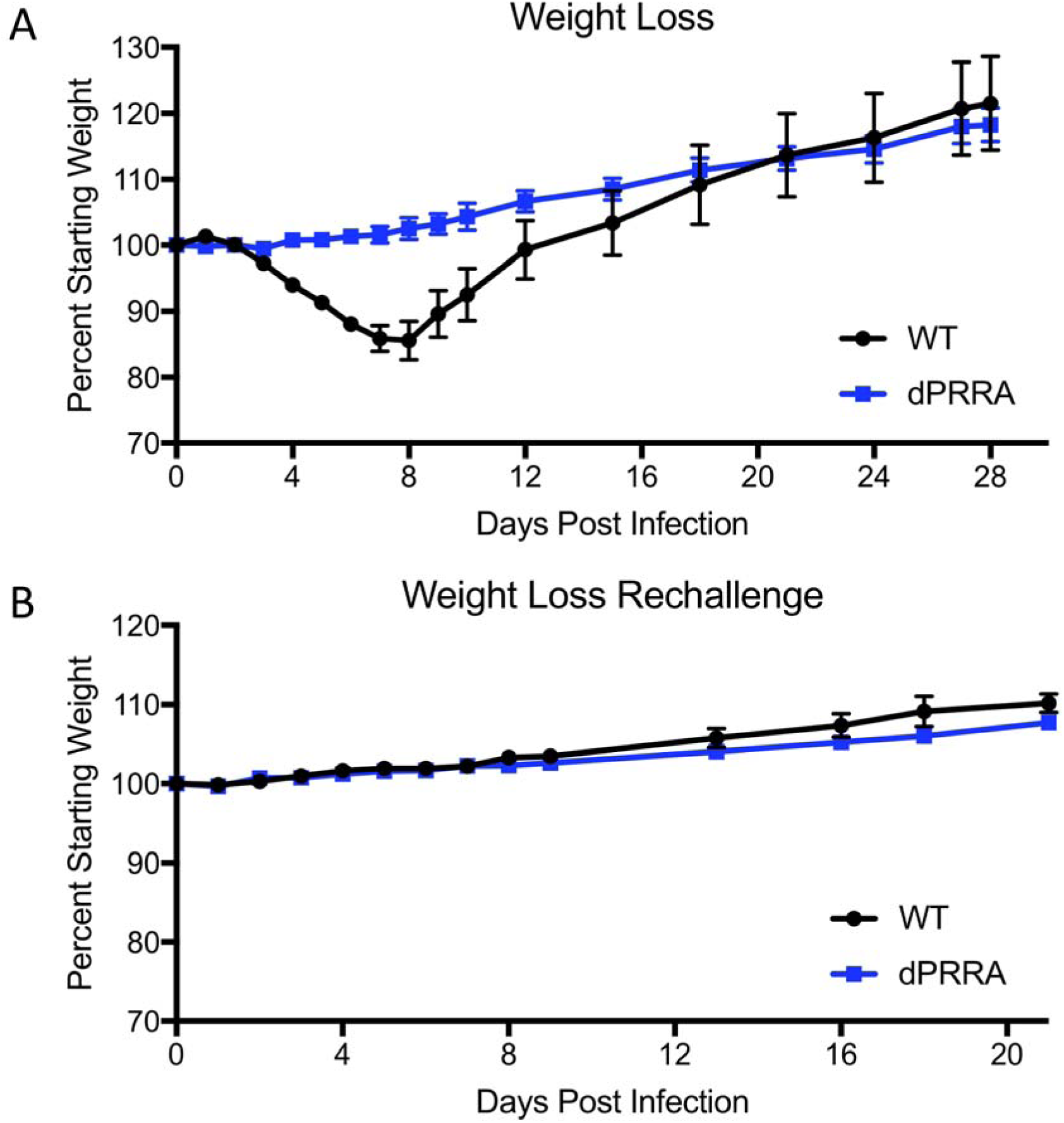
*In vivo* attenuation of ΔPRRA mutant. A) Weight loss following primary WT and ΔPRRA mutant SARS-CoV-2 challenge (N=4 per group). B) Weight loss following rechallenge of WT and ΔPRRA mutant infected mice with WT SARS-CoV-2 (N=4 per group).

